# Reinforcement learning-guided control strategies for CAR T-cell activation and expansion

**DOI:** 10.1101/2023.07.14.548968

**Authors:** Sakib Ferdous, Ibne Farabi Shihab, Ratul Chowdhury, Nigel F. Reuel

## Abstract

Reinforcement learning (RL), a subset of machine learning (ML), can potentially optimize and control biomanufacturing processes, such as improved production of therapeutic cells. Here, the process of CAR-T cell activation by antigen presenting beads and their subsequent expansion is formulated *in-silico*. The simulation is used as an environment to train RL-agents to dynamically control the number of beads in culture with the objective of maximizing the population of robust effector cells at the end of the culture. We make periodic decisions of incremental bead addition or complete removal. The simulation is designed to operate in OpenAI Gym which enables testing of different environments, cell types, agent algorithms and state-inputs to the RL-agent. Agent training is demonstrated with three different algorithms (PPO, A2C and DQN) each sampling three different state input types (tabular, image, mixed); PPO-tabular performs best for this simulation environment. Using this approach, training of the RL-agent on different cell types is demonstrated, resulting in unique control strategies for each type. Sensitivity to input noise (sensor performance), number of control step interventions, and advantage of pre-trained agents are also evaluated. Therefore, we present a general computational framework to maximize the population of robust effector cells in CAR-T cell therapy production.

**Author Summary:** Custom control strategies for expansion and activation of patient-specific CAR T-cell therapies resolved by reinforcement learning using a simulation environment and updatable cell growth parameters.

## Introduction

CAR T-cell therapy is a promising approach for personalized cancer treatment, applicable to a growing range of diseases such as treatment of B cell malignancies(1), multiple myeloma (2), solid tumors(3), HIV and several other types of cancer (4). In brief, CAR-T cell therapeutics involve production, collection, and separation of naïve T cells from the patient, transfecting them to produce Chimeric Antigen Receptors (CARs), and expanding them to provide a suitable dose. The cells are then infused back into the patient where they efficiently attack the malignant cells (5). During CAR T-cell therapy production, an important step is to activate the T cells, as they proliferate more rapidly than naïve, such that CARs are more readily expressed(6). One popular approach to activate the cells is by using antigen presenting beads, however prolonged proximity to aAPCs can lead to cell exhaustion (7–10). Exhausted cells consequently lose reproductive and therapeutic capacity (11). The success of an activation and expansion campaign is to have the maximum number of robust active cells. Logically, there is an optimal strategy to activation (bead addition) such that maximum cells remain activated, while the number of exhausted cells is minimized(6). However, such optimal strategies are often confounded by non-isotropic activation and propagation rates of donor cells based on age and other genetic factors(12,13). We thereby posit that the addition of activating beads is a dynamic control problem that must adapt based on the specific patient cell features and thus, is a good candidate for real time control strategies.

The current practice for activating T-cells is to add beads at the beginning of the culture and remove at the end (7,14,15). Prolonged signaling causes exhaustion and this can be mitigated by halting expression when unnecessary (9). As younger cells likely corelate to higher proliferation rates, the manufacturing time is often curtailed *ad hoc* (16). Although it is observed that intermittent exposure to bead yields a greater number of robust effector cells, the underlying activation-exhaustion mechanism, and its response to dosing across all cell types remains elusive till date (17,18). There is room for improvement considering the difficulty of the production process (7). No monitoring or control is involved in the activation process, which could partially explain the loss of potency of manufactured CAR T-cells (19). Optimizing CAR-T production is not limited to activation bead timing; cytokines, such as IL2, are necessary for survival and growth but also can induce Fas-mediated AICD (apoptosis induced cell death)(20). Treatment efficacy can be enhanced by controlling the dosage of IL-27 (21) while excessive IL-2 concentration leads to exhaustion (15). While the complex and rapid interplay between different cytokines and metabolic and genetic pathways in individual cells is hard to map, tracking the sensor outputs (22),(23) and imaging data (24) from the bulk population on-the-fly is far more amenable.

Recently there has been significant progress in the field of real time cell monitoring and automated control of biological processes (25). It is possible to track, monitor and infer the conditions of the individual cells from morphology (26). Sensors can output high dimensional feature space which can be used to profile cells or infer growth trajectories (27). Learning based algorithms (28) can be applied to navigate the decision making of adding/removing the activator. Reinforcement learning (29), is well adapted to solve such black-box decision problems where the system dynamics do not obey defined analytical expressions.

RL comprises an agent which interacts with an environment and gleans rules to develop a control policy (Figure 1a). At each time step, the agent assesses the current environment (observation space) and takes an action from a predefined list of allowed moves. The environment then updates based on the action and a reward or penalty is assigned to the agent which then updates their policy based on this learning. By repeated interaction with the environment, the agent continuously updates the learned strategy and establishes a policy. The policy is a function that maps every possible observation state to an action with a goal of maximizing the end reward. RL has been widely used for chatbots (30), autonomous vehicles (31), robot automation (32), predicting stock prices and projections (33), and industrial processes such as manufacturing and supply chain (34). Agents can perform better in an actual environment after being trained on incrementally complex simulated environments (35). RL algorithms are divided into two types – model based and model free. Model free RL algorithms optimize a policy or value function instead of modeling the environment. It can learn directly from sensor data and is useful in situations where it is difficult to model the environment. Despite being a well-established field, application of RL to optimization biological systems (36) is largely untapped. The main reason can be attributed to lack of suitable environments to train the agent and the confounding, inherent variability in biological processes. To benchmark new RL algorithms, OpenAI has established a test platform called gym(37), which has several environments on which new policy algorithms can be tested. The greatest benefit of using an RL platform is easy integration of different algorithms and policies to test on the specific environment. There are different environments coded for specific control tasks, for example - for simple robotic task – robo-gym(38), multi goal robotic task - panda-gym(39) and gym-pybullet-drones(40) and for self-driving bots – MACAD-gym (41). Biological processes are different than existing control problems like driving a car considering the complexity and variability. Each action by the agent on biological ‘environment’ will produce a stochastic outcome rather than a deterministic one. Controlling biological systems through RL will be possible with better understanding of system dynamics as well as design of better process simulations.

**Figure 1:**
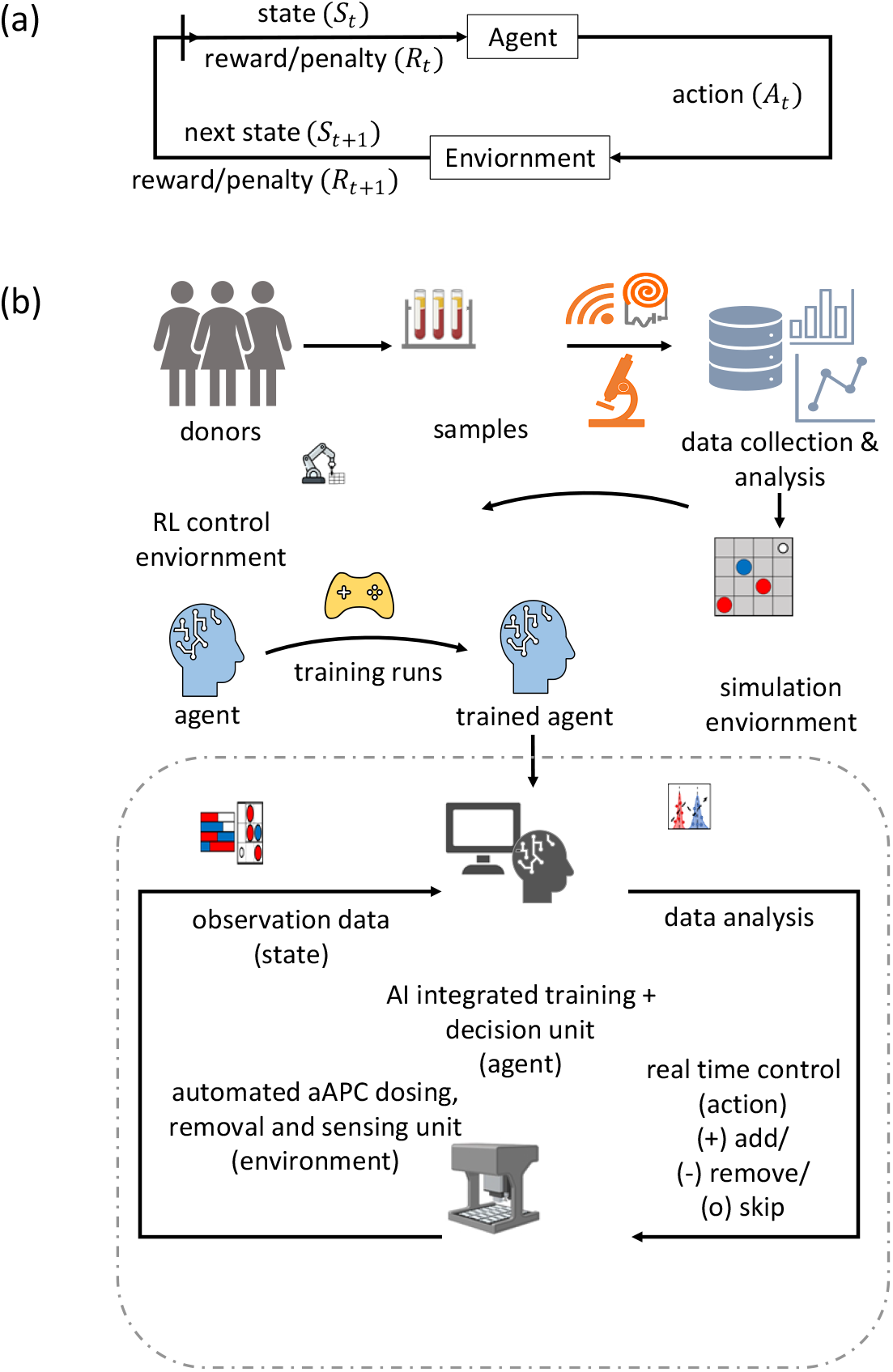
Reinforcement learning framework; (a) basic RL loop (b) RL workflow applied to real time control of T cell activation and expansion. The cell profiles and properties are inferred from donor sample pool with the help of imaging and sensing instruments. These properties coupled with data driven approach are used to create a simulation of the cell culture process. Afterwards this environment is transformed into the form of a game or RL control environment. An agent is trained on RL environment to get trained and navigate the actual cell activation process.

Multiple efforts have been made thus far in modeling T-cell expansion (42). Researchers have presented defined, analytical models with systems of ordinary differential equations (43). For modeling biological systems, stochastic models are often better suited than deterministic models. For instance, Monte Carlo (44) methods have been used to model CD4+ T-cells response to infection (45) and to manage biological variability for cell therapy production (46). Growth of organisms can be modelled with lattice kinetic MC simulation (47), for example Hall et al. modelled growth of yeast under influence of nutrient concentration and magnetic field exposure (48). Agent based models (ABM) is another stochastic approach where each component of the model is an autonomous entity governed by its own rules. ABM is widely used in T cell therapy models; for instance, Neve-Oz et al. presented agent-based simulations of T Cell -aAPC interactions (49) and Azarov et al. modeled chemotaxis of T-cell to dendritic cell (50).Zheng et al. have demonstrated a hybrid-RL strategy to optimize media replacement steps in cell therapy production and show *via* simulation that it outperforms deterministic models (51). Although control of the real physical environment (Figure 1b) is the near-term goal, demonstrating RL on a simulated biological process is a good first step to understand how RL can be effectively applied. Overall, the emergence of faster computing architecture is propelling us to a future of ML-driven policy making and training robotic arms for precision medicine and bioengineering.

In this paper, CAR T-cell activation and expansion is first coded as a 2D simulation, where a player or agent-algorithm can decide to add, skip, or remove antigen presenting beads to a given population of T cells in each step with the end goal of getting the highest number of potent cells when the simulation ends. This simulation can be termed as an agent-based model because each cell acts as an agent and behaves with a predefined set of rules (note, this use of agent is different from the RL agent discussed prior). The simulation is then converted into a customized gym environment in OpenAI Gym which enables testing several RL algorithms to benchmark policies for this custom environment. An RL agent algorithm then settles on an optimized strategy by repeatedly interacting with the environment. Three model free algorithms –proximal policy optimization (PPO)(52), actor-critic algorithm (A2C)(53) and deep Q-learning network (DQN)(54) are selected as candidate algorithms and are trained in this environment using three different observation spaces: 1) list of cell counts and other measurable parameters, 2) image of 2D cell environment, and 3) a combined list-and-image approach (Supplement 5). Different cell types are then used to test how the policies adapt their control strategies of bead dosing. The effect of noise resulting from poor sensors on training efficiency is also tested with observation variables corrupted with gaussian noise. Finally, the effects of changing the number of times the agent is allowed to interact with the environment as well as effects of pre-training agents on control performance are also tested and discussed.

## Results

### Design of CAR T-cell Activation Simulation

The objective of the simulation is to maximize the number of activated CAR T-cells through dynamic control of bead addition and removal. At each time interval, simulated data of culture condition in the form of tabulated sensor measurements and/or microscopic images will be provided to the agent. The agent can either add more beads, take away all beads or refrain from acting at that step (Figure 1b). After sufficient training in a similar environment, the agent is expected to choose an action based on the input data which will optimize the end goal. Before attempting such a control strategy on a physical environment, the process of bead-based CAR T-cell activation is simulated as an RL environment (Figure 1a). A 2D surface (Figure 2c) for cell growth is simulated as a continuous *n* × *n* grid with spacing of 10 microns to match the approximate cell diameter (55). In all the simulation 50×50 corresponding to 500 by 500 sq-micron area is used. For better clarity in observing the cells, (in Figure 2c) a 20 × 20 grid is used for demonstration purposes. The simulated expansion area is made continuous (no boundary). All defined parameters for this simulation are described in Table 2. Although attempts were made to associate these parameters with literature values, some assumptions were made. It is important to note that the modular simulation and RL training presented here can be readily updated as more measured values are determined. A fixed time method (56) is used with a value of 6 min per step, derived from the approximate time a cell translates one diameter away or to the next grid spacing (velocity of the cell is ∼2 micron per minute (50)). There are other factors affecting cellular migration like media viscosity, age of cell, size of cell, etc. that are neglected in this simplified model. The total simulation lasts for a 7-day expansion campaign, equivalent to 1600 simulation steps. Bead to cell contact, bead to cell ratio and confluence are taken into consideration in the simulation rules considering their role in the efficiency of activation (57).

**Figure 2:**
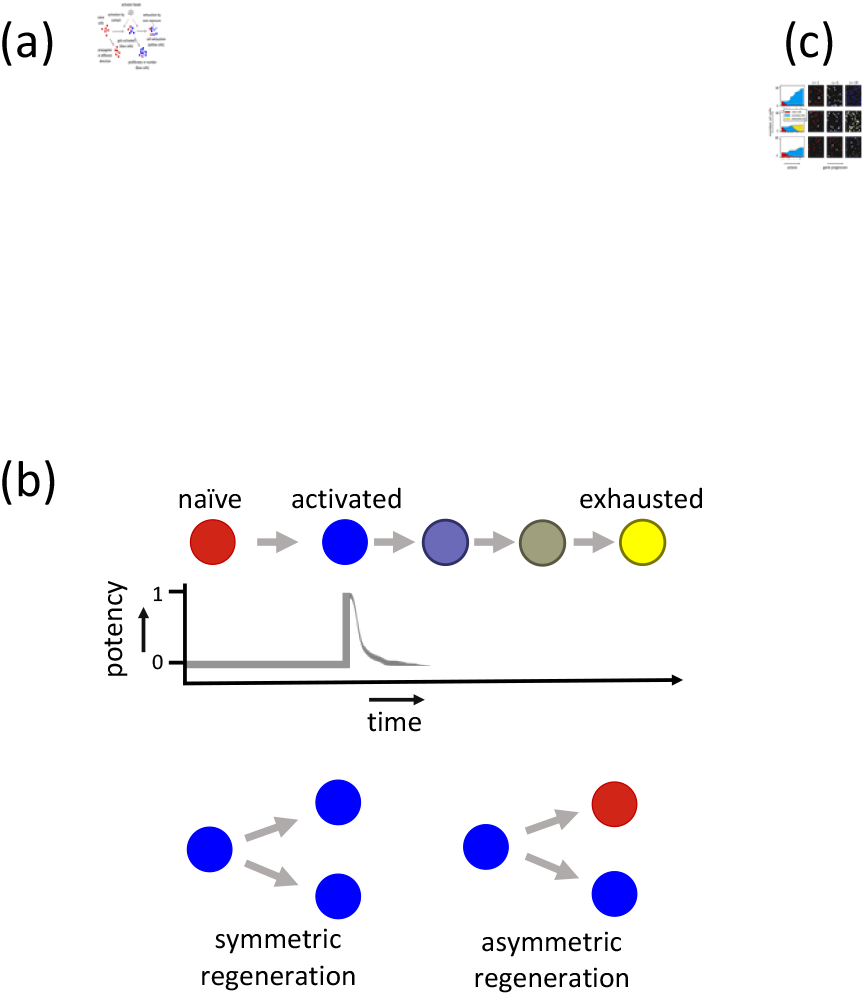
Proposed simulation replicating cell activation and expansion (a) Process and permitted actions by the cells in each simulated step. (b) Simulated life trajectory of a naïve starting cell to activated with full potency, and natural exhaustion caused by aging. Also defined are two modes of division – symmetric and asymmetric. (c) Sample simulation trajectories for three control strategies – top to bottom row depicts optimum, sub-optimum and random bead additions; bar plot at left indicates the number of cells separated by type at each simulation step, the symbols at the x axis represents the action taken: (+) refers to bead addition, (−) refers to removal and (o) refers to no action; the right three windows are simulation screens at 1, 5 and 19 steps.

At simulation start the grid is randomly seeded (Figure 2c) with a specified number of naïve T-cells indicated as red cells in the simulation. The following steps are iterated for each cell in the simulation: *Step 1.* It can propagate to any of the 8 adjacent cells if it satisfies movement conditions, namely vacancy at the chosen grid and probability of making a move at that step determined stochastically (Figure 2a and Supplement 2). *Step 2:* If a naïve cell occupies a position where an activation bead (coupled to anti-CD3 and anti-CD28 antibodies) is present and if certain conditions (probability of conversion at that step beyond a threshold determined stochastically, detailed in Supplement 2) are met, the naïve cell is activated and turns blue in the simulation (Figure 2a). *Step 3:* If an already activated cell gets in a position where there is a bead, it gets exhausted depending on the value of the specified exhaustion rate (Figure 2a). *Step 4:* At each time step each activated cell gets exhausted as per natural, transient exhaustion rate (58) which is 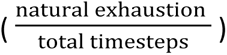 times smaller compared to accelerated exhaustion caused by over exposure and stimulation caused by beads (see Table 1 Units and Figure 2b).

**Table 1.**
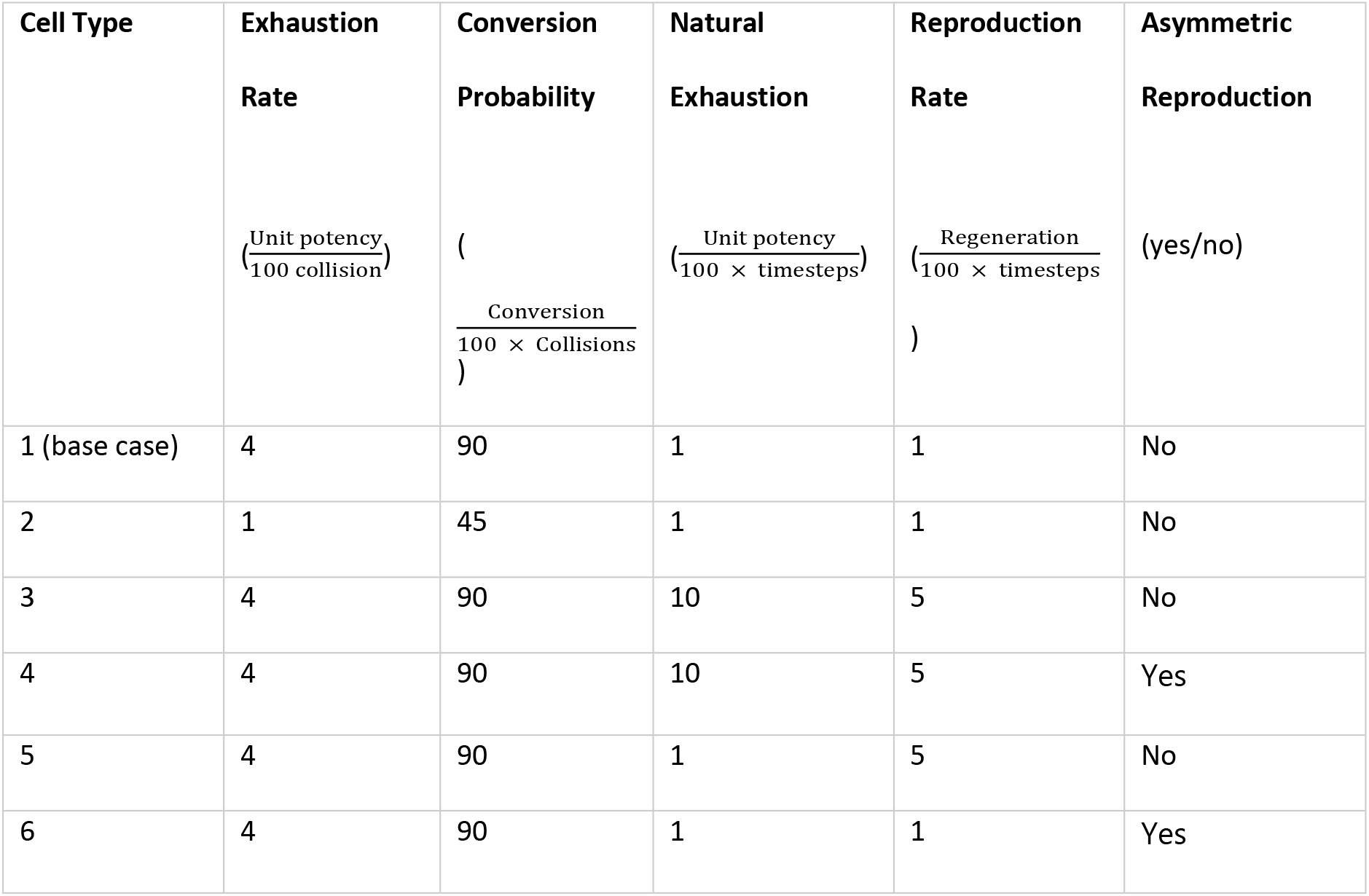
Simulated Cell Types.

| Cell Type | Exhaustion<br>Rate<br>$\left(\frac{\text{Unit potency}}{100 \text{ collision}}\right)$ | Conversion<br>Probability<br>$\left(\frac{\text{Conversion}}{100 \times \text{Collisions}}\right)$ | Natural<br>Exhaustion<br>$\left(\frac{\text{Unit potency}}{100 \times \text{timesteps}}\right)$ | Reproduction<br>Rate<br>$\left(\frac{\text{Regeneration}}{100 \times \text{timesteps}}\right)$ | Asymmetric<br>Reproduction<br>(yes/no) |
| --- | --- | --- | --- | --- | --- |
| 1 (base case) | 4 | 90 | 1 | 1 | No |
| 2 | 1 | 45 | 1 | 1 | No |
| 3 | 4 | 90 | 10 | 5 | No |
| 4 | 4 | 90 | 10 | 5 | Yes |
| 5 | 4 | 90 | 1 | 5 | No |
| 6 | 4 | 90 | 1 | 1 | Yes |

**Table 2:** Parameters and their descriptions.

| Variable Name | Value | Relevant Source |
| --- | --- | --- |
| **Simulation Variable** |  |  |
| Initial number of beads | 0 | (18) |
| Grid Number | 2500 |  |
| Grid dimension | 10 Micron |  |
| Confluence | Half of total grid (1250) | (57) |
| Number of control-steps | 20, 50, 400 (variable) | Assumed |
| Control Time interval | 8-hour,<br>3.2-hour ,<br>24 minutes | (18) |
| Number of beads that can be added at each control step | 10 | (18,68,69) |
| Initial number of naïve cells to begin with in control area | 20 | Assumed |
| Total time | 160 hours (weeklong growth) | (7) |
| **Cell Variable** |  |  |
| Mean value of Regeneration Age | 2 days | Assumed |
| Maximum age at which a cell can regenerate | 3.5 days | Assumed |
| Probability of conversion | $45, 90 \left( \frac{\text{Conversion}}{100 \times \text{Collisions}} \right)$ | Assumed |
| Exhaustion Rate | $1, 4 \left( \frac{\text{Unit potency}}{100 \text{ collision}} \right)$ | Assumed |
| Natural Exhaustion | $1, 10 \left( \frac{\text{Unit potency}}{100 \times \text{timesteps}} \right)$ | Assumed |
| Regeneration Rate | $1, 5 \left( \frac{\text{Regeneration}}{100 \times \text{timesteps}} \right)$ | Assumed |
| Asymmetric regeneration | True/False | Assumed |
| Potency value above which a cell is considered robust | 0.8 | Assumed |
| Potency value below which a cell is considered exhausted | 0.2 | Assumed |

Each cell has several attributes that are tracked through the simulation, such as activated potency which starts at 0 with naïve cells and steps to value of one when activated (Figure 2b). *Step 5:* An activated cell can proliferate under condition of matured age, potency, and stochastic probability (Figure 2a and detailed in Supplement 2).

The control agent can add beads, take out beads or skip taking any action at the time step. A set number of beads are added in each control step (e.g., 10, 20, 40). It is observed that optimum bead to cell ration vary widely from 3:1 to 9:1 depending on bead and cell type (59,60). Keeping this in consideration, as the system is seeded with 50 cells, 10 beads are allowed to be added in each control step (beads can be added in consecutive steps). If a control step occurs every 3.2 hours, there are 32 simulation steps between actions and the agent can take a total of 50 control actions for each simulation 1600/32). As an action the agent can either add, remove, or keep beads the same. In the removal step all the beads are taken out at once by a magnet (assuming use of commercial paramagnetic beads). This is one important real-world constraint where the agent does not have the choice to incrementally add or take out beads, it must add in a specified amount or take out all together at a single step. Based on the properties of cell (such as regeneration rate, how much it exhausts over time, chance of getting converted if encountering a bead) the sequence of action chosen by the agent can be optimal (ensuring high throughput of robust effector cells marked with bright blue), or sub-optimal (low number of effector cells or low potency effector cells marked with yellow) at the end of expansion steps (Figure 2c).

### Evaluating input strategies and algorithms

At each step of an RL episode, the agent algorithm chooses the action by taking an observation snapshot as input. There are many possible observation data formats that can be sent to the agent. For example, bulk measurements could be made by impedimetric (22) (Agilent, Xcelligence) or permittivity-based (61) sensors (Skroot Lab Inc). Real time imaging systems (23) (Sartorius Incucyte) coupled to Artificial Intelligence (AI) empowered cell classification tools can specify and quantify cell types based on morphology (62). Those tools can be used to count naïve and activated cells along with other cell properties such as age and robustness. Other data such as time elapsed, quantity of beads in the system and action history can be obtained from the system itself. All the data can be input in the form of a list of measured values to the agent. This method is termed the tabular method in this work (Figure 3). Another possible observation format can be in the form of image, obtained from high precision microscopy. In this work we also try to observe if a three-channel image of the simulation environment like Figure 2c alone is enough to provide the agent with enough information to adequately train (Figure 3). The third input format tested is the fusion between the above two, where both tabular and image information are provided to the agent (Figure 3). Here we refer to each agent in ‘algorithm-input’ format, for example PPO-image refers agent trained with PPO algorithm on image data. The aim of this analysis is to demonstrate how agent training depends on algorithms and input schemes.

**Figure 3.**
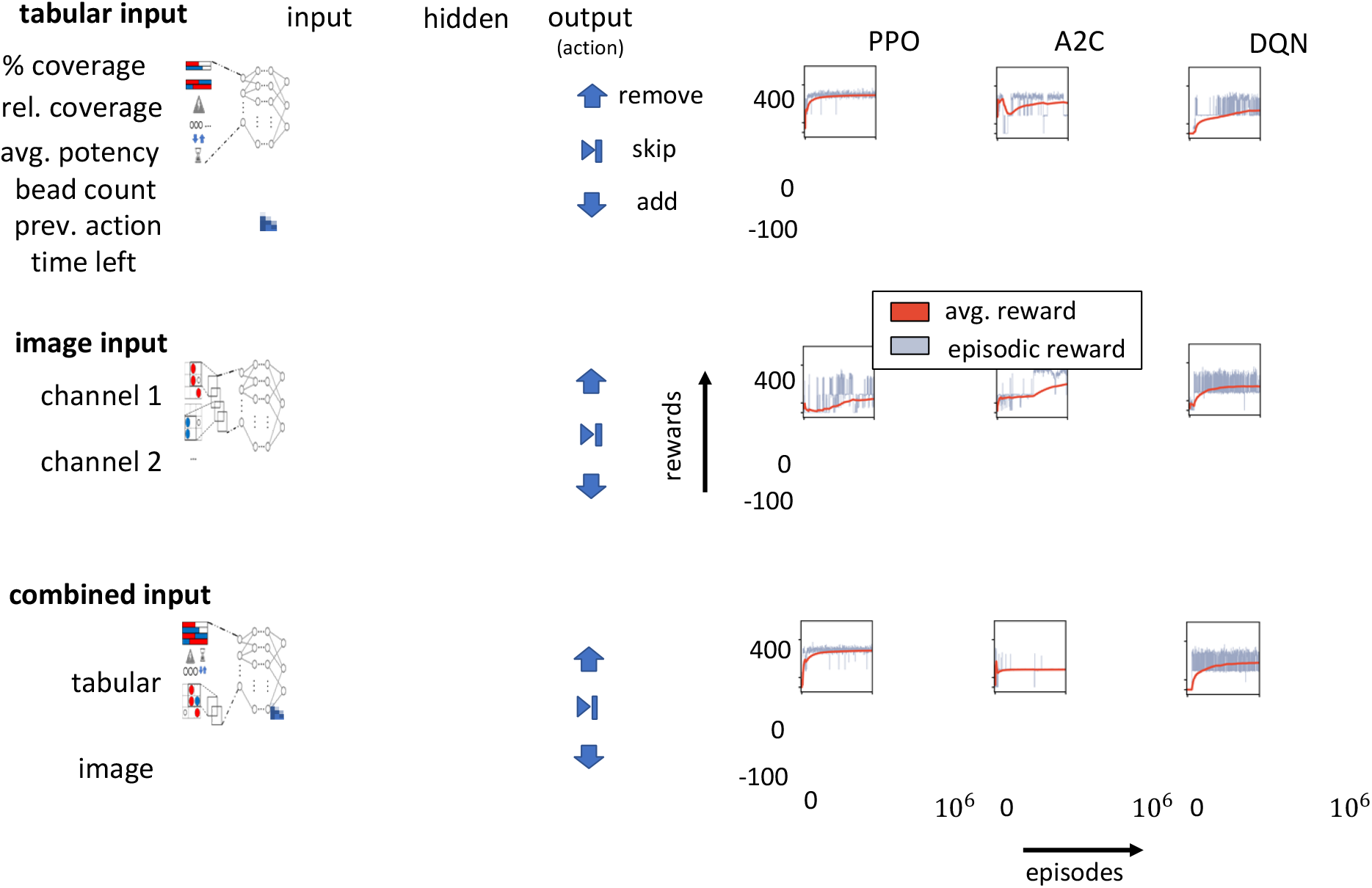
A higher average reward with tight distribution of outcomes are indications of a better trained agent. The quality of the policy can be determined from the episodic reward distribution of a trained agent. For example, with PPO-tabular and DQN-image (Figure 3), the agent adopted a stable strategy by 100,000 training episodes as observed from the episodic reward and flattened out average reward. But with PPO-tabular the episodic reward distribution around average is +/−50 whereas it is +/−250 for DQN-image. That indicates PPO-tabular agent is better trained which has a tighter distribution of higher rewards and DQN-image is subjected to variability and chance events. The distribution is even tighter for A2C-combined, but the average reward is far less than PPO-combined or DQN-combined.

The reward the agent gets at the end of each episode is the episodic reward and at each point we plot another point averaging all previous episodic rewards (average reward shown in red in Figure 3). The rising trend of the average reward in the beginning indicates the agent is learning and constantly obtaining a better strategy whereas flattening of the average reward indicates that the agent has settled for an optimized strategy (see PPO-tabular and DQN-tabular input in With image input, our goal is to observe if it is possible to navigate the environment by getting a snapshot of number of cells and beads, cell type, potency, and age from an image only without any temporal labels. We probed whether the simulation strategy can be step independent. We notice that context information is important. Performance for all algorithms were higher with context data (tabular and combined) than without-context data (image only). In all three-input strategies the nature of DQN was very similar. It settles for a sub-optimal strategy with broader reward distribution (details in *Discussion*). In this work the default hyperparameters for each neural architecture (Supplement 3,4,5) as reported in OpenAI Gym were used without fine-tuning. How an untrained and trained agent navigates the environment is demonstrated in supplementary video 1 and 2 respectively.

### Learned control strategies for different cell types and number of controls steps

Next, a PPO-combined agent is tested on each respective cell type, simulating diversity of patient-derived cells, to assess how the RL agent can adapt its learned control strategy. Six cell types are simulated by changing the cell parameters (Table 1). For each of the cell types, an agent is first trained for 1M timesteps and then used to navigate 1000 simulations on the same ‘environment’. The average number of beads in each control step is plotted with standard deviations to reveal the bead addition patterns (Figure 4). The variable actions taken in response to observations (presence of error bars) indicates that the policy is adaptive to navigate different situations and did not simply memorize and repeat the same actions at each step. In a few instances, there was uniformity of actions (no error bars, same number of beads in all 1000 simulations). The learning curve is also included with the bar plot (insets) indicating that the agent settled for a policy at the end of training (discussed above).

**Figure 4:**
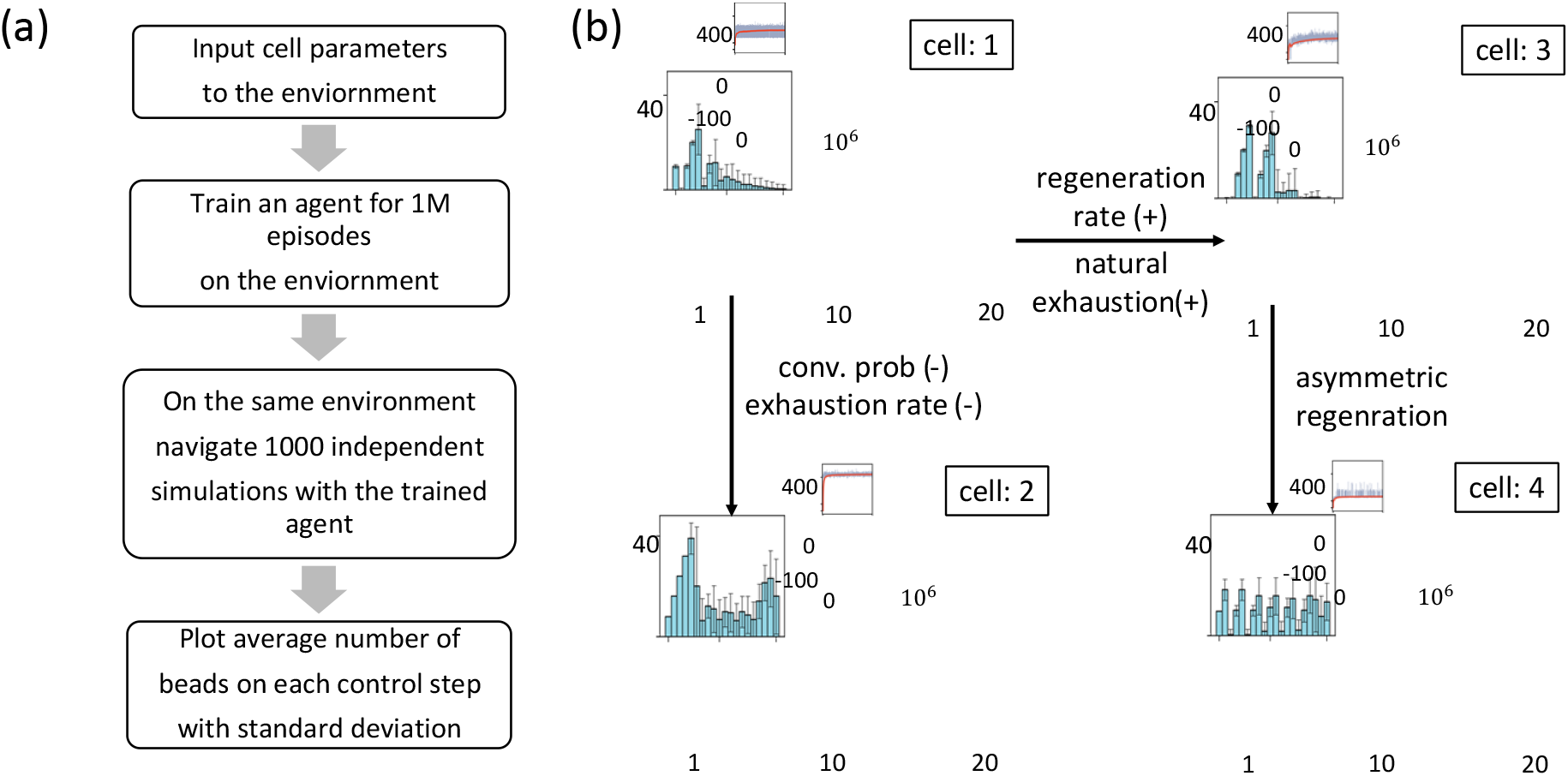
Change of strategy by the agent using 20 control steps for different cell types. (a) Simulation process to obtain control strategy information (b) Strategy of the agent visualized by average number of beads at each control step (y and x axes respectively). Error bar indicates the standard deviation of beads used at that control step – indication of simulation variability or constancy (where there are no bars). The learning curve is also attached with each bar plot, axes same as Figure 3. Arrows between plots indicate the change in cell type (also see Table 1).

The learned control strategies correlate with intuition for these extreme edge cases. In the base case of Cell 1, to protect the cells from overexposure it removes the beads on the second step after adding on the first. The intuitive strategy would be to add the beads in the initial steps and let most of the naïve cells convert and remove the beads when most cells are activated and let them proliferate and increase in number which is what the agent executes with less beads after step 5. With Cell type 2, that has lower rate of exhaustion than the base case, we observe the agent ramps up number of beads quicker and maintains a near constant level of exposure until the end when there is another ramp to activate any remaining naïve cells (Figure 4b). It is interesting to note that in this case, the first steps of the agent (the initial ramp) are very decisive, with no deviation amongst all runs. Afterwards there are variations in bead number with agent taking decision as required to convert the remaining naïve cells. In cell type 3, we simulate a cell that has a higher rate of natural exhaustion. As exhaustion is only applicable to active cells, the obvious strategy would be to make some deliberate delay in adding the beads, to convert the cells close to the end of the episode. However, as regeneration will be high the whole region will be crowded with activated cells so it would be imperative to remove beads and wait for all of them to regenerate as soon as the optimal number of cells get activated. Considering both cases the best strategy would be to add beads in the middle steps and skip the beginning and end steps. This is reflected in the learned strategy of the agent, it skips the first two steps, then adds the beads in two repeated steps, then takes out all the beads and waits to make the cells increase in number. With cell type 4, asymmetric regeneration is simulated where an activated cell can produce both activated and naïve cells.

To convert the newly produced naïve cells, beads are required, but those same beads cause the activated cells to get exhausted. To navigate this system the agent alternately adds and removes beads, and the overall end score is lower than the other cell types.

To test the effect of an agent that has more control over the environment, we repeat the training process with 50 control steps (interacting with the growth vessel every 3.2 hr instead of 8 hr – see justification in Supplement 8) for six cell types (Table 1). The base case behaved the same way, with more dosing of beads in the beginning and reduced in the end (Figure 5). But as it has more frequent control points, the agent skips adding beads at the onset to account for small natural exhaustion, continuously adding beads for second to fifth step, then performed the add-remove-skip step depending on simulated status, with diminishing number of beads in subsequent steps. For cell type 2, it adds beads for more steps at the outset (Figure 5) than before (Figure 4b) and Cell types 3 and 4 differ as well. Cell 5 is simulated with only regeneration increased from the base case and the agent removes beads in the second half to let the activated cells grow without getting exhausted. However, when the natural exhaustion is increased in cell type 3, the agent falls into a dilemma: if it adds bead at the beginning, the converted cells will be exhausted in the next steps, if it adds bead at the end, it cannot take advantage of the higher regeneration rate. Balancing these constraints, the agent adds beads in the first two steps and then removes them in the third and skips the next 10 steps. It then adds or removes bead depending upon the present situation. However, this is less favorable than other cell types and ends with a lower number of potent cells in the end. Finally, for cell type 6 we increased the rate of natural exhaustion and added asymmetric regeneration. In this case the agent alternately adds and removes beads for first third of the control steps, and then ramps number of beads with variability based on current cell count; again, the expected outcome (average reward) for this unfortunate cell type is dependent on chance and lower than others.

**Figure 5:**
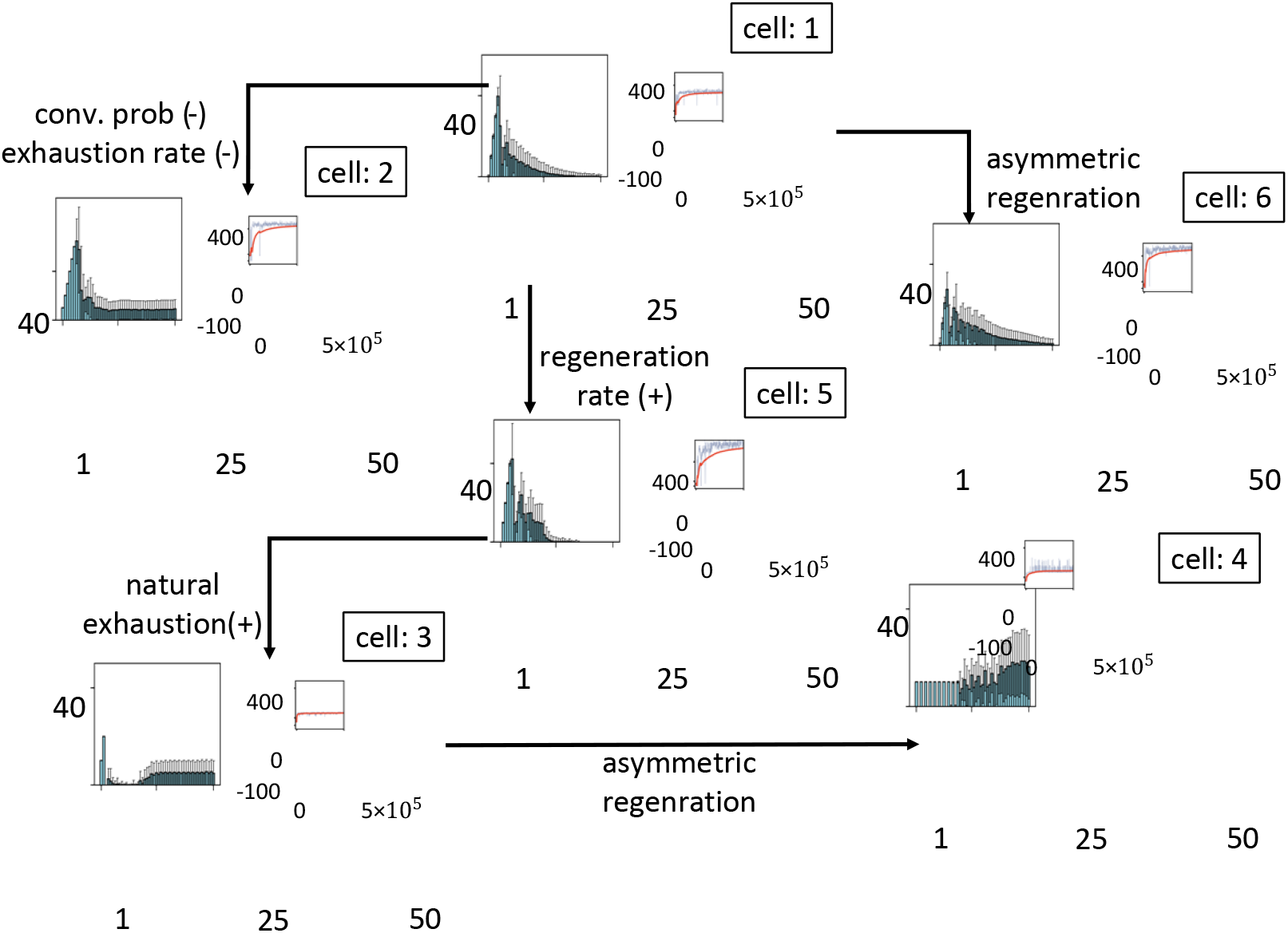
Change of strategy by the agent using 50 control steps learned from training with different cell types. Strategy of the agent visualized by average number of beads per control step (y and x axes respectively). Error bars indicate one standard deviation, showing variability of steps or uniformity (no error bars). The learning curve is also attached with each bar plot. Arrows indicate the change in cell type, also see Table 1.

### Effect of measurement noise, number of control steps, and number of training runs

The ability of an agent to learn unique control strategies for different cell types is a major finding; however, to put this into practice, it will be important to know how accurate the measurements (inputs to agent) must be as well as the required number of training runs (as 10^6^ experiments to determine a unique training regime is not tractable). Here we explore both topics using the T-cell expansion simulator using the PPO algorithm with combined input.

The observation space for tabular input would be obtained from cell monitoring sensors that distinguish between cell types and estimate potency (optical, impedance, etc.). These devices will not have complete precision. To observe the effect of noise, an agent is trained with 40% of the initial cell number added as gaussian noise in cell count and potency estimation to simulate measurement error. There is no observable change in the episodic and average reward of the training steps and reward distribution with and without noise (Figure 7a). There are two possible reasons: first, gaussian noise in a stochastic environment does not make perceivable difference in mapping observation to action and second, the agent either maps the noise along with the observations or totally disregards the noisy observations and build its policy on more stable inputs such as time steps. A histogram is also drawn at three stages of training – the zeroth training run, where the agent is fully random, as well as at 250k and at 500k episodes. It is also observed that there is a clear difference in the reward distribution between the random agent at start and trained agent at 250k runs, but the distribution of rewards at 250k and 500k episodes were indistinguishable.

Figures 4 and 5 demonstrate that the agent can perform better with increased interaction with the environment (50 control steps rather than 20). With more interaction it has better control and there is a higher reward with less fluctuation whereas with fewer interactions it is difficult to control the environment. We investigated if this pattern holds for even further interactions.

Conceivably, an agent could interact with a fully automated environment at every observation point. To observe the effect of increased control, we trained an agent with 400 control steps (adding, removing, or maintaining beads every 24 m). In this case there are an overwhelming 3^400^ possible combinations of action sequences. With such a high number the agent finds it difficult to settle on a control policy and the learning curve fluctuates more than the 50-control point case (Figure 7b). This finding indicates that ‘real-time’ control is likely not as advantageous as a control strategy that is still dynamic yet has a tractable number of possible actions.

In a realized, clinical setting, there will likely be a limited number of experiments that can be performed on a new cell type (patient sample) for the agent to self-learn a control strategy. The average learning curve of cell 1 shows 90% of max average reward after 29,000 training sessions for an agent with 50 control steps (Figure 6c). We hypothesized that this number could be further reduced if an agent trained on one cell type is then used as start point for another cell (e.g., training the agent on a stock cell, prior to testing with the patient cell sample). To test this approach, the agent is trained on 500k training runs on a base case cell 1 and then used to subsequently train on Cell types 1-4. For cell 1 and cell 2 the optimum strategy is similar – to add beads in the beginning. In that case the agent can adapt faster, and a smaller number of runs (1000 or one updated policy step) is required in comparison to training from scratch to reach the same level of accuracy. But optimum strategy is different for cell 3 and 4 – to add beads at the end. In those cases, the agent needs to unlearn the previous strategy and adapt a new strategy. With such a change in policy, it takes longer to reach the same level of accuracy rather than starting training from scratch. An alternative or parallel approach to settling on an optimal control strategy would be taking patient cells and performing a series of tests to obtain growth parameters that would allow for efficient simulation (Figure 1b). Then *in silico* tests, much like this, would augment the physical training data. An *in-silico* test thus can guide if there is a change in policy and weight the choice of – retraining on another cell or training from scratch considering desired yield and resources.

**Figure 6:**
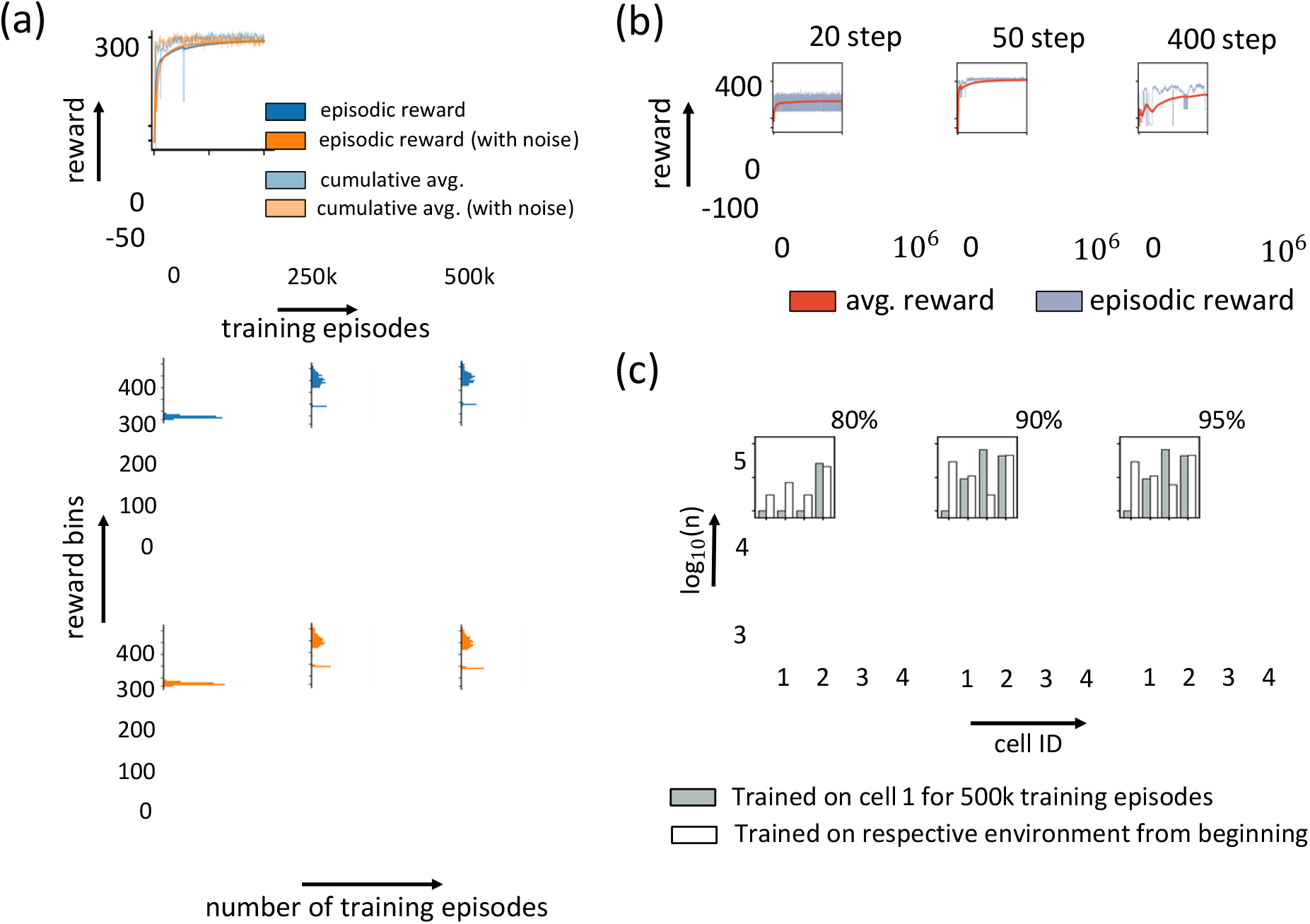
(a) Learning curve for agent trained with and without noise and reward histogram for simulation conducted with agent trained on 0, 250k and 500k episodes (b) Agent trained with 20, 50 and 400 timesteps (c) Number or training episodes required to reach accuracy of 80%, 90% and 95% by agents pre-trained for 500k steps on cell 1 vs. agents trained on respective cell types from beginning. Y axis shows the number of training runs required in log base 10 scale.

## Discussion

Here we elucidate a possible RL-based platform which would help robotic arms to precisely deploy or remove activator molecules at specific time points during T-cell activation to ensure maximum number of activated cells (i.e., peak therapeutic potential) prior to pushing them back to the patient. As a steppingstone, cell growth parameters were directly obtained and inferred from literature to simulate the spatial and temporal stochasticity of CAR T-cell activation and expansion with reasonable fidelity. These simulation parameters, like tuning knobs, can be updated to reflect accurate metrics of cell growth in a plug-and-play fashion for a given experiment. Before deploying this neural engine (agent) for a specific experiment, we first pre-train it to understand properties of the cell type at hand and its physiochemical attributes. This should reduce the number of training runs required in the physical environment. The platform also pinpoints key patient-specific humoral parameters that should be measured and imputed to an *in-silico* model – thereby making this relevant for personalized, point-of-care therapeutics. The observables (physical parameters or snapshot of a lattice of cells) from the physical environment collected as sensor output or imaging data, will be continuously fed as input to RL to device the best dosing policy to maximize activated cells. Continued research on accurate, non-invasive, real-time measurement techniques to enumerate cell types during culture will provide faster training performance. With an improved and realistic simulation and adequate data on patient-specific cell parameters, one could adjust the simulation parameters (Table 1) and perform pre-training of the control strategy *in silico* before coupling to the physical environment. The simulation would consequently inform the type of sensors needed for the physical environment. It would also show how much noise the agent can accommodate before it fails to learn anything at all. With a large amount of measurement noise, the agent will likely (a) disregard the noisy observation parameters (e.g., cell number, cell type, and potency), and (b) fix a redundant policy based only on simulation step count.

Notwithstanding, one possible reason for the agent’s ineptitude to learn solely from a discrete image input (Figure 3) owes to the lack of connection with the preceding and succeeding time-points. Thus, it becomes next to impossible to gauge whether a certain action (dosing) indeed helped in maximizing the number of robust cells. To this end, we hope that instead of just providing one disembodied frame, if we exposed the model to short stacks of three to five consecutive frames, the learning rate and gains might improve – but we leave this as an exercise for the future.

Moving forward, this cell-activation routine guided by RL can be used as a template for other model-free, stochastic biological applications. Apart from CAR T-cell activation, this bears promise to unravel hitherto unknown biological policies opted by nature such as – the underlying optimization of cell differentiation and cell proliferation. With the advent of modern generative algorithms and data driven approaches, we can hypothesize that it will be possible to create digital twins of cell culture environments. Additionally, this 2D simulation can be updated to a 3D environment representing more realistic growth conditions in static reactors (multilayer growth). Possible further experiments are enlisted on Supplement 8. In addition, this work provides a basis to benchmark Transformer (63), and DAL-e (64) based implementations of the same, which are finding increasing applicability in different domains of biology. A library of such pre-trained models enacted by robotic arms for precision dosing would be useful to match the range of cell types handled by clinics of tomorrow. This will enable effective control of activation and expansion and get more efficacious therapies to patients faster.

## Methods

### Simulation Design

The simulator of cell expansion was made using the Pygame (65) module of python and is hosted on Zenodo (66) and GitHub - https://github.com/Sakib1418/Game-of-cells. The simulation was designed to integrate with OpenAI gym (37), a collection of simulated environments and associated toolkits to test and compare RL agent algorithms. As the new gym environment was made, the Stable Baselines3 module (67) was used on top of gym to explore current RL algorithms. The properties of the actors (cells) attempt to simulate actual CAR T-cells, for example movement and regeneration rate. Due to current lack of measured parameters such as conversion probability on encountering a bead, reasonable estimates are made in this initial work. All simulation values and cell parameters are listed in Table 2. To observe agent response with different cells, new cell types are conceptualized by changing these cell-properties (Table 1). How these parameters are formed into equations governing the fate of the cell and on the the culture environment or simulation trajectory overall is detailed in the game pseudocode (Supplement 2). Installation of the simulation-game, data analysis and reproduction of the plots and usage are detailed in the project GitHub repository.

### Reward Function Design

In RL, the agent gets a reward or penalty at each step dependent on its latest action. The purpose of a good reward function is to direct the agent towards its desired outcome as efficiently as possible. Designing a reward function is an iterative process that considers an understanding of the algorithms and environment. For this cell activation and expansion control task, a reward scheme is proposed in which the end goal is to get the greatest number of potent activated cells. For this we must encourage the agent to add beads to activate the cell as well as remove beads when it estimates exhaustion. At each step if the average potency is higher than the previous step the agent receives a small reward of 5, otherwise it receives a penalty. At the last step the sum of potency of all the cells above a threshold is multiplied by 100 and added to the reward. The reasoning behind this scheme is that at the initial part of the training the small rewards in all the timesteps encourage the agent to add beads to activate the cells, because this way the average potency per cell increases.

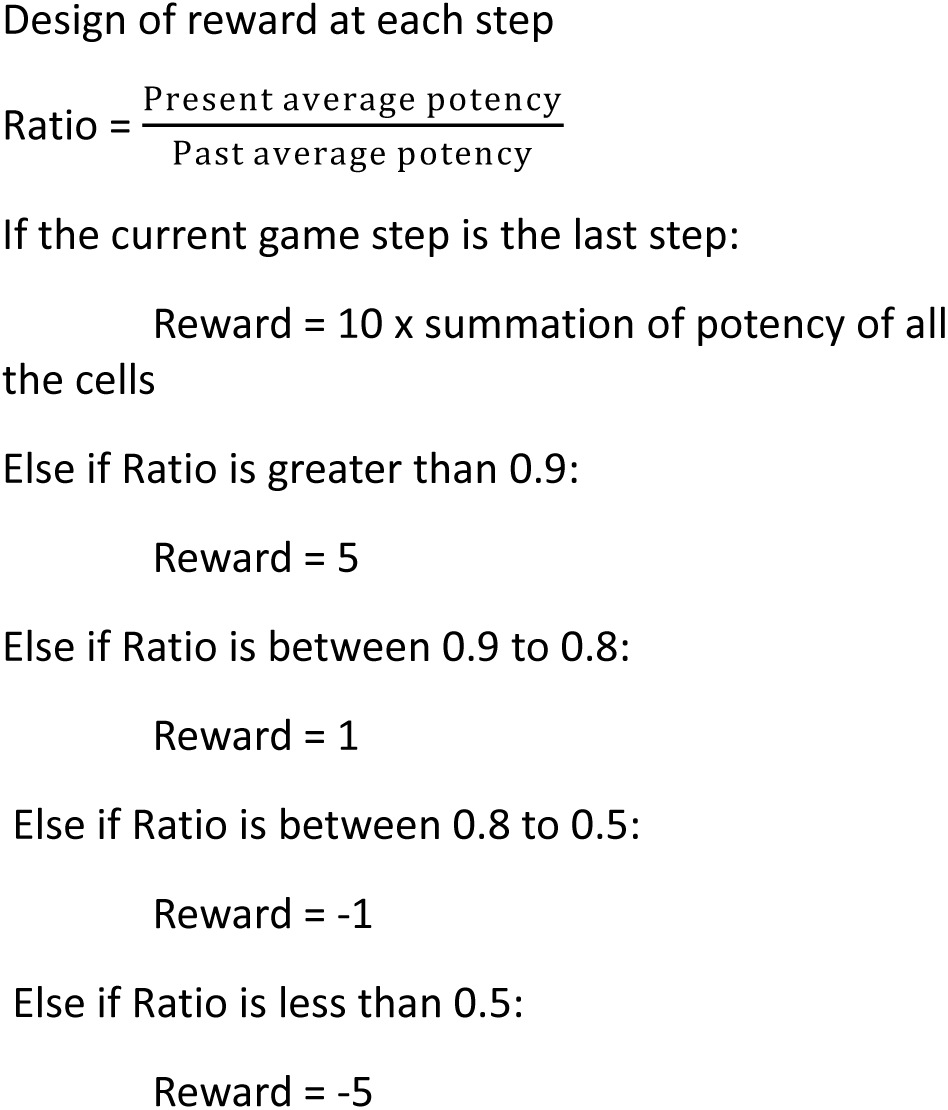

After a few steps, the existing beads have the chance to cause exhaustion to activated cells The reward obtained for the end goal is much higher than the rest of the timesteps combined, which prompts the agent to take steps to score higher at the end even at the expense of sacrificing some initial rewards. In this way, by repeated interaction with the environment an agent can self-train and determine when to add or take out beads to maximize the reward score. Apparently, it seems the ratio value higher above one should be rewarded more, but empirically we found out that the agent falls at reward frustration and fails to learn if higher values are used at the beginning.

## Acknowledgements

We thank Krishanu Saha and his group members for useful discussions on CAR-T cell activation and expansion.

## Author Contribution

SF performed the literature review, conceptualization, data acquisition, code, and software compilation, writing and reviewing of the manuscript. IFS assisted in code compilation and version control. RC contributed to writing, supervision, and revision. NFR contributed to conceptualization, funding acquisition, writing, reviewing and overall supervision.

## Correspondence

All the correspondence should be addressed to Nigel Forrest Reuel at

## Code Availability

Code is available in Zenodo at - https://doi.org/10.5281/zenodo.7905320 and at GitHub at - https://github.com/Sakib1418/Game-of-cells with full details and instructions for reproduction.

## Supplementary Material

### Policy Networks

**Table 1:**
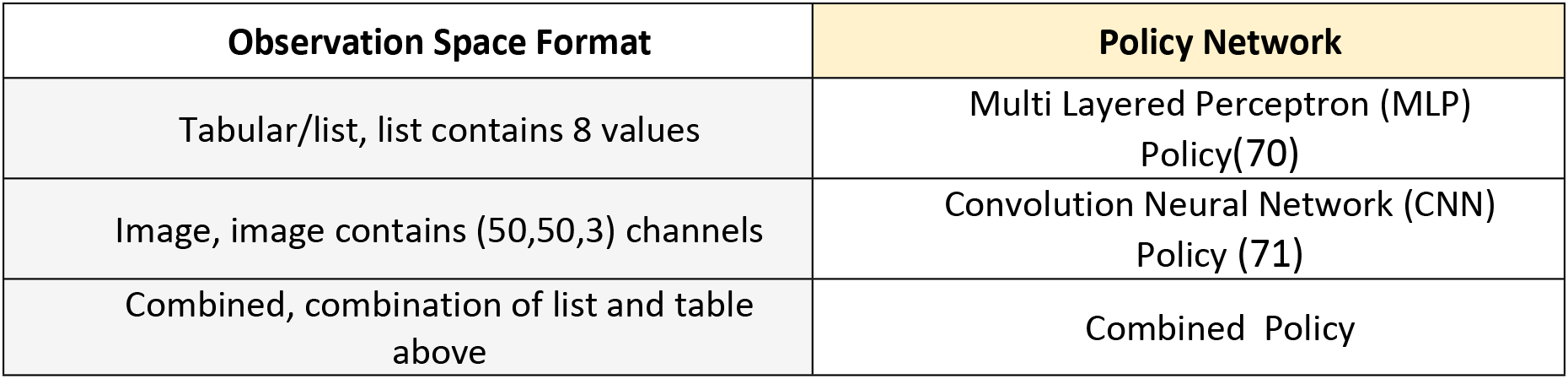
Policy Network used for tabular, image and combined inputs. Detail can be found at this link.

| Observation Space Format | Policy Network |
| --- | --- |
| Tabular/list, list contains 8 values | Multi Layered Perceptron (MLP)<br>Policy(70) |
| Image, image contains (50,50,3) channels | Convolution Neural Network (CNN)<br>Policy (71) |
| Combined, combination of list and table<br>above | Combined Policy |

### Simulation Pseudocode

The game pseudocode is given below. The bold words indicate function name. The function description follows the pseudocode.

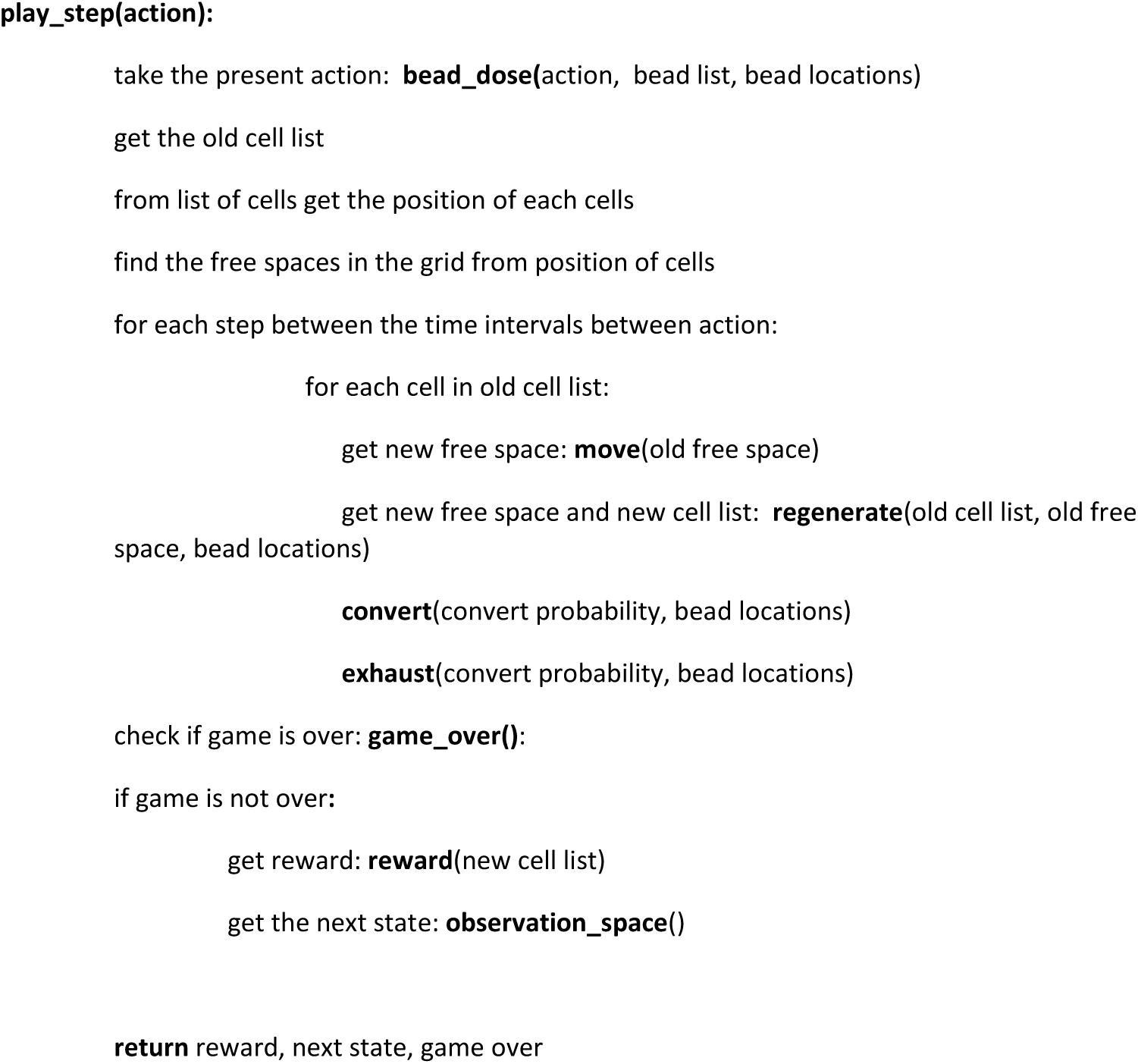

The play_step is called by the agent at each step of the game. Several functions are used to carry out the tasks. Each of them is described.

#### bead_dose()

This function works based on 3 actions –

If the action is to add the beads it drops 10 beads in random positions in the grid.

If the action is to remove beads it takes out all the beads

If the action is to skip it makes no change to the present bead number

**Figure 1:**
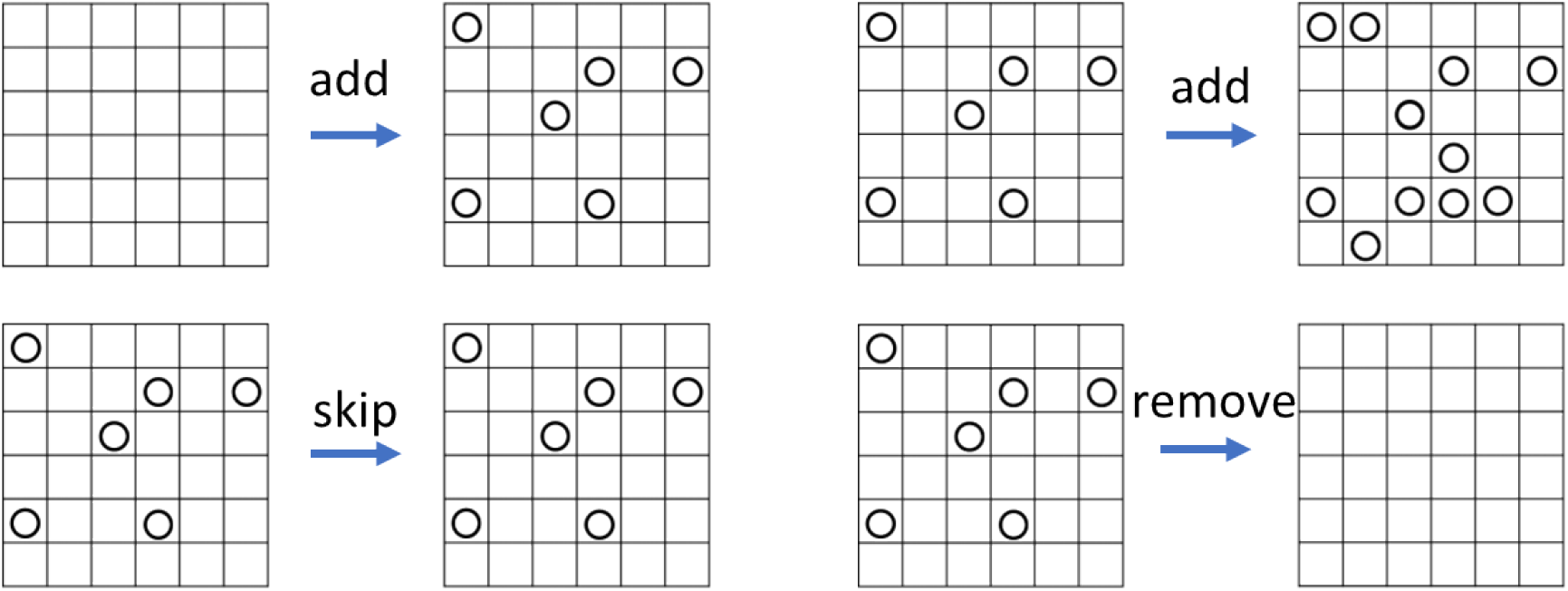
Agent action involving aAPC bead dosing demonstration with 6 by 6 grid space with 5 beads

#### move()

Moves the present cell into account into a new cell if all these conditions are met–

It passes a probability based on some predefined distribution (tunable from variable.py) above a threshold. Then it chooses any of the 8 grids around it.

If the grid of its choice is empty at that step.

This function returns the new free space after the move.

**Figure 2:**
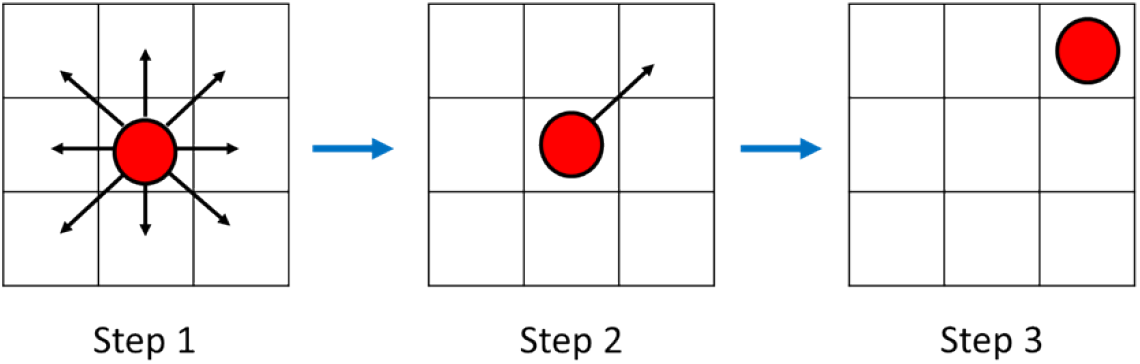
Steps in cell movement - step 1: A cell have possible 8 directions to move to its adjacent grid, step 2: It selects a direction and checks for the criteria mentioned above, step 3: It moves to that grid.

#### regenerate()

A new cell is created adjacent to the previous cell if all these following conditions are met – If the present cell into account is in reproductive age (defined in cell parameters at table 2) of main text and supplement 3.

If the chosen grid is vacant.

If the cell is potent

If the cell is active

If a probability threshold (tunable from variable.py) is passed

**Figure 3:**
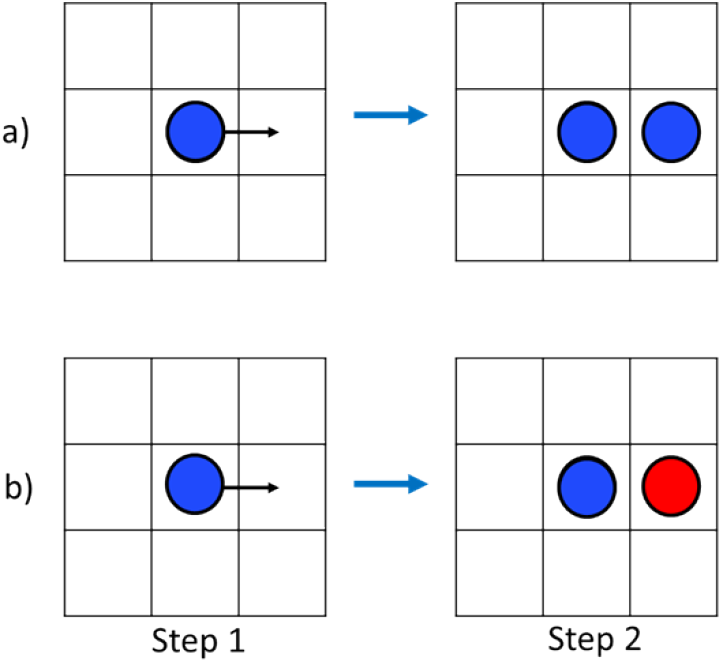
regeneration steps - step1: the cell selects a direction step 2: if all the stated conditions are fulfilled it produces another activated or naive cell at that grid

#### convert()

Converts a red(naïve) cell to a blue(activated) cell if all these following conditions are met – The cell is naïve.

A bead is in the same position as the cell.

A probability threshold is passed (name of the variable is convert probability, tunable from variables.py script)

**Figure 4:**
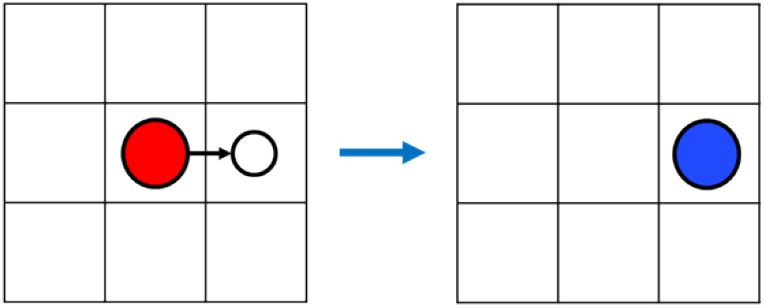
Cell differentiation to activated cell, if a naive cell goes to a grid with bead and conditions fulfilled the cell differentiates to an activated cell.

#### exhaust()

The potency of the cell is reduced by exhaustion rate if –

The cell is activated.

A bead is in the same position as the cell.

The cell is exhausted by an amount much smaller than exhaustion rate in all other cases.

**Figure 5:**
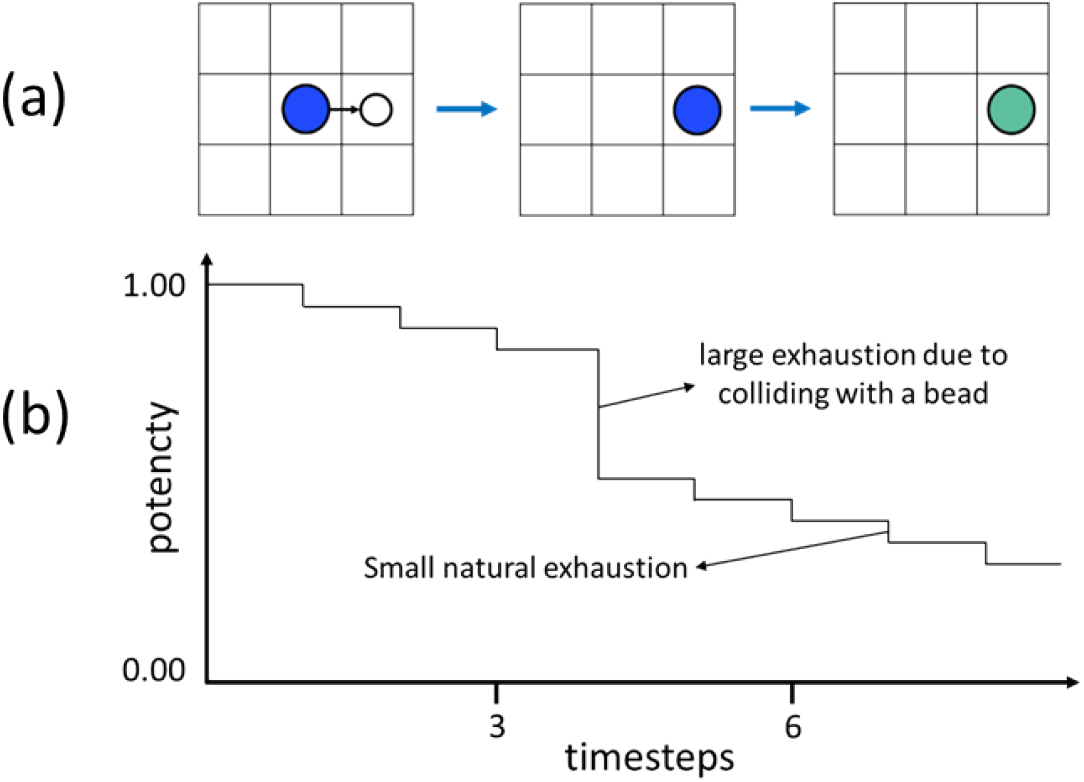
(a) exhaustion procedure (b) potency loss over timestep (not scaled)

#### game_over()

Returns a Boolean – True if any one of the criteria is met –

The simulation is run more than the maximum timestep.

A certain portion of the grid (tunable) is covered by the cells.

#### reward()

In all steps except the last assigns a small reward depending on the average potency of the cells. In the last step if the average potency is more than a certain threshold, assigns a big reward on that step.

#### observation_space()

Depending on the observation space type, it returns an image or a list. The list contains – fractional coverage of grid by active and naïve cells, relative coverage by active and naïve cells, number of beads, total potency of the cells, previous action and time elapsed. The image contains – the 3-channel image of the simulation display.

### Algorithm Structure: Tabular observation space

~~~
 ActorCriticPolicy(
  (features_extractor): FlattenExtractor(
   (flatten): Flatten(start_dim=1, end_dim=-1)
  )
  (pi_features_extractor): FlattenExtractor(
   (flatten): Flatten(start_dim=1, end_dim=-1)
  )
  (vf_features_extractor): FlattenExtractor(
   (flatten): Flatten(start_dim=1, end_dim=-1)
  )
  (mlp_extractor): MlpExtractor(
   (shared_net): Sequential()
   (policy_net): Sequential(
    (0): Linear(in_features=8, out_features=64, bias=True)
    (1): Tanh()
    (2): Linear(in_features=64, out_features=64, bias=True)
    (3): Tanh()
   )
   (value_net): Sequential(
    (0): Linear(in_features=8, out_features=64, bias=True)
    (1): Tanh()
    (2): Linear(in_features=64, out_features=64, bias=True)
    (3): Tanh()
   )
  )
  (action_net): Linear(in_features=64, out_features=3, bias=True)
  (value_net): Linear(in_features=64, out_features=1, bias=True)
 )
~~~

### Algorithm Structure: Image observation space

~~~
 ActorCriticCnnPolicy(
  (features_extractor): NatureCNN(
   (cnn): Sequential(
    (0): Conv2d(3, 32, kernel_size=(8, 8), stride=(4, 4))
    (1): ReLU()
    (2): Conv2d(32, 64, kernel_size=(4, 4), stride=(2, 2))
    (3): ReLU()
    (4): Conv2d(64, 64, kernel_size=(3, 3), stride=(1, 1))
    (5): ReLU()
    (6): Flatten(start_dim=1, end_dim=-1)
   )
   (linear): Sequential(
    (0): Linear(in_features=256, out_features=512, bias=True)
    (1): ReLU()
   )
  )
  (pi_features_extractor): NatureCNN(
   (cnn): Sequential(
    (0): Conv2d(3, 32, kernel_size=(8, 8), stride=(4, 4))
    (1): ReLU()
    (2): Conv2d(32, 64, kernel_size=(4, 4), stride=(2, 2))
    (3): ReLU()
    (4): Conv2d(64, 64, kernel_size=(3, 3), stride=(1, 1))
    (5): ReLU()
    (6): Flatten(start_dim=1, end_dim=-1)
   )
   (linear): Sequential(
    (0): Linear(in_features=256, out_features=512, bias=True)
    (1): ReLU()
   )
  )
  (vf_features_extractor): NatureCNN(
   (cnn): Sequential(
    (0): Conv2d(3, 32, kernel_size=(8, 8), stride=(4, 4))
    (1): ReLU()
    (2): Conv2d(32, 64, kernel_size=(4, 4), stride=(2, 2))
    (3): ReLU()
    (4): Conv2d(64, 64, kernel_size=(3, 3), stride=(1, 1))
    (5): ReLU()
    (6): Flatten(start_dim=1, end_dim=-1)
   )
   (linear): Sequential(
    (0): Linear(in_features=256, out_features=512, bias=True)
    (1): ReLU()
   )
  )
  (mlp_extractor): MlpExtractor(
   (shared_net): Sequential()
   (policy_net): Sequential()
   (value_net): Sequential()
  )
  (action_net): Linear(in_features=512, out_features=3, bias=True)
  (value_net): Linear(in_features=512, out_features=1, bias=True)
 )
~~~

### Algorithm Structure: Combined observation space

~~~
MultiInputActorCriticPolicy(
(features_extractor): CombinedExtractor(
(extractors): ModuleDict(
(image): NatureCNN(
(cnn): Sequential(
(0): Conv2d(3, 32, kernel_size=(8, 8), stride=(4, 4))
(1): ReLU()
(2): Conv2d(32, 64, kernel_size=(4, 4), stride=(2, 2))
(3): ReLU()
(4): Conv2d(64, 64, kernel_size=(3, 3), stride=(1, 1))
(5): ReLU()
(6): Flatten(start_dim=1, end_dim=-1)
)
(linear): Sequential(
(0): Linear(in_features=256, out_features=256, bias=True)
(1): ReLU()
)
)
(vector): Flatten(start_dim=1, end_dim=-1)
)
)
(pi_features_extractor): CombinedExtractor(
(extractors): ModuleDict(
(image): NatureCNN(
(cnn): Sequential(
(0): Conv2d(3, 32, kernel_size=(8, 8), stride=(4, 4))
(1): ReLU()
(2): Conv2d(32, 64, kernel_size=(4, 4), stride=(2, 2))
(3): ReLU()
(4): Conv2d(64, 64, kernel_size=(3, 3), stride=(1, 1))
(5): ReLU()
(6): Flatten(start_dim=1, end_dim=-1)
)
(linear): Sequential(
(0): Linear(in_features=256, out_features=256, bias=True)
(1): ReLU()
)
)
(vector): Flatten(start_dim=1, end_dim=-1)
)
)
(vf_features_extractor): CombinedExtractor(
(extractors): ModuleDict(
(image): NatureCNN(
(cnn): Sequential(
(0): Conv2d(3, 32, kernel_size=(8, 8), stride=(4, 4))
(1): ReLU()
(2): Conv2d(32, 64, kernel_size=(4, 4), stride=(2, 2))
(3): ReLU()
(4): Conv2d(64, 64, kernel_size=(3, 3), stride=(1, 1))
(5): ReLU()
(6): Flatten(start_dim=1, end_dim=-1)
)
(linear): Sequential(
(0): Linear(in_features=256, out_features=256, bias=True)
(1): ReLU()
)
)
(vector): Flatten(start_dim=1, end_dim=-1)
)
)
(mlp_extractor): MlpExtractor(
(shared_net): Sequential()
(policy_net): Sequential(
(0): Linear(in_features=264, out_features=64, bias=True)
(1): Tanh()
(2): Linear(in_features=64, out_features=64, bias=True)
(3): Tanh()
)
(value_net): Sequential(
(0): Linear(in_features=264, out_features=64, bias=True)
(1): Tanh()
(2): Linear(in_features=64, out_features=64, bias=True)
(3): Tanh()
)
)
(action_net): Linear(in_features=64, out_features=3, bias=True)
(value_net): Linear(in_features=64, out_features=1, bias=True)
)
~~~

### Algorithm Analysis

Among the three algorithms discussed, PPO (72) and A2C (53) are policy optimization algorithms and DQN belongs to Q learning algorithms. PPO optimizes a surrogate function, A2C performs a gradient ascent and DQN (54) learns an approximator function (73). PPO and A2C are policy algorithms which means the update uses data from the latest policy. DQN is an off policy where each update is independent of policy.

DQN is subject to variability as seen from all input types (Figure 3), due to many failure modes. Relative performance of PPO was better as it has several advantages over the other methods, such as -sample efficiency and moderate policy updating using a clipped surrogate function. The probable cause of A2C failing in all modes may be its sensitivity to non-stationary stochastic environment and hyper parameter choice.

### Time-step Justification

The total simulation consists of 1600 smallest time unit of 6 minutes. If the agent intervenes 50 times, then between each intervention there will be 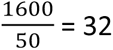 steps or 32 × 6 = 192 minutes or 3.2 hours. Similarly, if the agent intervenes 20 times, then between each intervention there will be 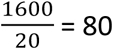 steps or 80 × 6 = 480 minutes or 8 hours.

### Possible Future Experiments

There are numerous possible further experiments that could be performed using the proposed environment for CAR T-cell activation. A non-exhaustive list of the experiments is given below –

In this work the built in hyperparameters and policy algorithms (details in Supplement 1) found in OpenAI Gym are used to test the simulation and algorithms. Researchers can find experimentally derived values (table 2) and design better simulations, customized policy network and neural network architecture with tuned hyperparameters. We postulate that in this way, it is possible to come up with an optimized strategy for cell expansion, but this is beyond the scope of this work and can be a good follow up study.

Alternate to solely retraining on a physical environment, cell parameters can be obtained from patient-sample tests to update a custom *in silico* environment. With simulations it is possible to provide a greater number of training runs, however the efficacy of this training is dependent upon 1) how accurately cell parameters can be obtained and 2) how true to real-world the simulation is. Simulations also provide the opportunity to explore even more fine-tuned process control – such as inclusion of spatiotemporal control.

In this initial work, beads were dosed in a fixed quantity and when withdrawn all the beads were removed. One modification would be to train the agent to dose/withdraw beads at specific places and at a range of quantities. In these cases, spatial data (image characterization) as well as good feature extraction algorithms will be required.

The applicability of a trained model on a new cell type can be estimated with simulation; if the policy needs to be revised completely, like the agent trained on cell 1 and then applied to cell 4, the model must be restarted and trained anew.

Apart from custom *in silico* environment, generative algorithms can be used to produce realistic cell culture/activation data as observation space and train the learning agent.

